# Redefining the Game: MVAE-DFDPnet’s Low-Dimensional Embeddings for Superior Drug-Protein Interaction Predictions

**DOI:** 10.1101/2024.04.01.587541

**Authors:** Liang-Yong Xia, Yu Wu, Longfei Zhao, Leying Chen, Shiyi Zhang, Mengdi Wang, Jie Luo

## Abstract

Precisely predicting drug-protein interactions (DPIs) is pivotal for drug discovery and advancing precision medicine. A significant challenge in this domain is the high-dimensional and heterogeneous data characterizing drug and protein attributes, along with their intricate interactions. In our study, we introduce a novel deep learning architecture: the <u>M</u>ulti-view <u>V</u>ariational <u>A</u>uto-<u>E</u>ncoder embedded within a cascade <u>D</u>eep <u>F</u>orest (MVAE-DFDPnet). This framework adeptly learns ultra-low-dimensional embedding for drugs and proteins. Notably, our t-SNE analysis reveals that two-dimensional embedding can clearly define clusters corresponding to diverse drug classes and protein families. These ultra-low-dimensional embedding likely contribute to the enhanced robustness and generalizability of our MVAE-DFDPnet. Impressively, our model surpasses current leading methods on benchmark datasets, functioning in significantly reduced dimensional spaces. The model’s resilience is further evidenced by its sustained accuracy in predicting interactions involving novel drugs, proteins, and drug classes. Additionally, we have corroborated several newly identified DPIs with experimental evidence from the scientific literature. The code used to generate and analyze these results can be accessed from https://github.com/Macau-LYXia/MVAE-DFDPnet-V2.

## I. Introduction

Over the past few decades, a myriad of computational methodologies for the identification of DPIs have been devised, substantially narrowing the search scope for potential drug and protein candidates. This advancement has markedly diminished the costs and boosted the efficiency of drug discovery and development processes. Typically, these methodologies fall into three distinct classes: ligand-based [1], molecular docking [2], and machine learning-based methods [3]–[7]. While the ligand-based approach relies on a sufficient number of known ligands for a given protein; the Molecular docking approach is limited to available 3D protein structures [8]. Conversely, machine learning-based methods have emerged as a highly promising avenue for predicting DPIs [9]–[12].

Over the years, the input features for machine learning-based approaches have evolved from pharmacogenetics, which comprises information on drugs (i.e., chemical structures) and proteins (i.e., encoding sequences) represented as feature vectors [13], [14], to similarity network-based input features [15], and more recently, the trend has shifted toward incorporating heterogeneous networks that integrate various biological and pharmacological data [5], [16], [17]. Such heterogeneous data sources provide a rich and inherently related information, offering a multi-view perspective for the prediction of novel DPIs. On one hand, incorporating these diverse data sources can enrich the feature set and potentially boost prediction accuracy. On the other hand, it also increases input dimensionality, posing challenges for subsequent analyses.

The application of deep learning techniques further enhances the power of drug-protein pair prediction models. DeepWalk [18] built a tripartite, heterogeneous network from biomedical linked datasets and utilized the network’s node similarity for prediction. NeoDTI [17] used the neighborhood information of the network and learned topology preserving representations of drugs and proteins. deepDTnet [19] adopted a deep auto-encoder to learn high-quality features from heterogeneous networks and then applied positive-unlabeled matrix completion to predict new DPIs. AOPEDF [20] employed arbitrary-order proximity to derive a low-dimensional representation of drugs and proteins, and subsequently developed a cascade deep forest classifier to predict new interactions.

While these methods have each made notable strides in the field, there remains substantial room for improvement. A particularly challenging task lies in the construction of compact, low-dimensional embedding of drugs and proteins that are consistent across different types of drug/protein-related networks. The use of deep learning as a classifier is also challenging due to the need to fine-tune numerous hyperparameters. As such, the construction of low-dimensional drug and protein embedding, and the design of an effective deep learning-based pipeline are urgently required for efficient and accurate DPIs prediction.

To address these issues, we develop MVAE-DFDPnet, a network-based framework for DPIs prediction that fuses a multi-view variational auto-encoder (MVAE) with a cascade deep forest (CDF). The strengths of MVAE-DFDPnet are twofold: Firstly, it effectively consolidates individual networks into a unified low-dimensional embedding representation; secondly, it employs a deep cascade forest classifier, as described by Zhou et al. [21], which delivers high-performance classification with significantly fewer hyperparameters than traditional deep neural networks. This deep forest allows for automatic complexity adjustment. The synergy between advanced data compression and a flexible classification approach culminates in a robust, sophisticated, and appreciably enhanced model for drug discovery.

We evaluate MVAE-DFDPnet on benchmark datasets and compare its performance against several state-of-the-art methods. Our analysis reveals that MVAE-DFDPnet outperforms its counterparts in predictive accuracy, achieving this with a reduced number of drug and protein embedding. We also evaluate the robustness of MVAE-DFDPnet using previously unseen drug-protein pairs, demonstrating its high performance under stringent conditions. Finally, we provide visualizations of the learned drug and protein embedding and validate novel interactions predicted by our model.

## II. Materials and Methods

### A. Dataset

We integrated heterogeneous bio-networks as multi-view data inputs, encompassing four distinct entity types (Drugs, Proteins, Diseases, and Sideeffect) and 15 different types of associations; further details are provided in Table I. Our task was to infer unknown DPIs within a network comprising 732 Food and Drug Administration (FDA)-approved drugs and 1,915 unique proteins. Within this network, 4,978 known DPIs are labeled as positive samples, with a corresponding number of randomly-selected non-interacting (or ‘unknown’) pairs labeled as negative samples. For more details on dataset collection, readers are referred to the recent works by Luo et al. [5] and Zeng et al. [19].

**TABLE I:** Summary of heterogeneous biological networks as multi-view data.

| Category | View | Association | Drug | Protein | Disease | Sideeffect | No.of.Egde | Database |
| --- | --- | --- | --- | --- | --- | --- | --- | --- |
| Drug | 1 | Drug-Drug | 732 | - | - | - | 132768 | [22] |
|  | 2 | Drug-Disease | 732 | - | 440 | - | 1208 | [22], [23] |
|  | 3 | Drug-Sideeffect | 732 | - | - | 12904 | 263805 | [24] |
|  | 4 | Drug chemical similarity | 732 | - | - | - | 118578 | [25] |
|  | 5 | Drug therapeutic similarity | 732 | - | - | - | 240825 | [26] |
|  | 6 | Drug sequence similarity | 732 | - | - | - | 215902 | [22] |
|  | 7 | Biological Processes similarity | 732 | - | - | - | 243812 | [27] |
|  | 8 | Cellular Component similarity | 732 | - | - | - | 271192 | [27] |
|  | 9 | Molecular Function similarity | 732 | - | - | - | 244226 | [27] |
| Protein | 1 | Protein-Protein | - | 1915 | - | - | 112468 | [28] |
|  | 2 | Protein-Disease | - | 1915 | 440 | - | 23080 | [29] |
|  | 3 | Protein sequence similarity | - | 1915 | - | - | 1705532 | [30] |
|  | 4 | Biological Processes similarity | - | 1915 | - | - | 1611052 | [27] |
|  | 5 | Cellular Component similarity | - | 1915 | - | - | 1665635 | [27] |
|  | 6 | Molecular Function similarity | - | 1915 | - | - | 1653812 | [27] |

### B. Methods

In this paper, we propose a novel network-based method termed MVAE-DFDPnet, which achieves substantial feature compression along with a flexible prediction mechanism.

MVAE-DFDPnet takes multi-view data as input. The biological interaction network of each view is preprocessed into a probabilistic co-occurrence (PCO) matrix [19], then calculate a shifted positive pointwise mutual information (PPMI) matrix by following Bullinaria and Levy [31]. The next phase involves inputting a drug/protein’s high-dimensional feature vector into the MVAE to produce a reduced-dimensionality embedding. Lastly, a CDF is then employed to predict DPIs using the concatenated embedding of drug-protein pairs. The output is a binary indicator that denotes the presence or absence of an interaction between a specific drug-protein pair. We sequentially present the stages of our methodology in Fig. 1, progressing from left to right.

**Fig. 1:**
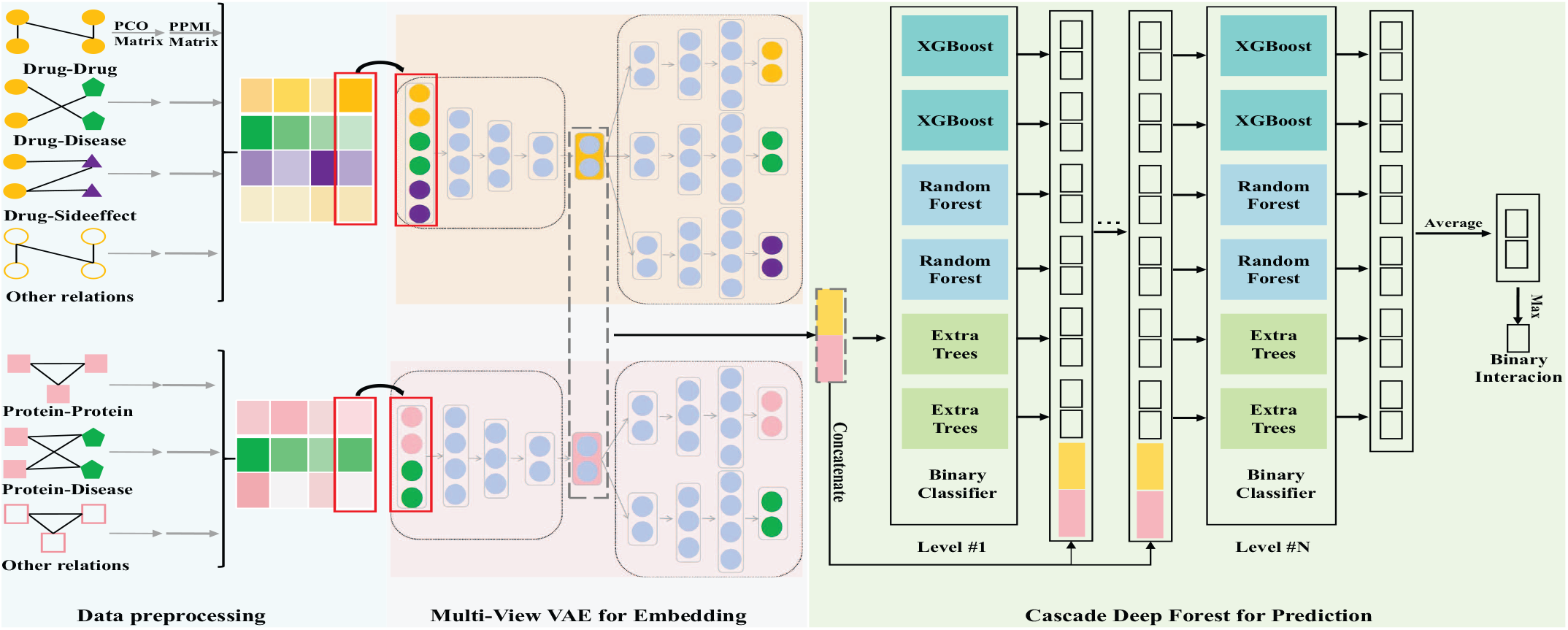
The schematic flowchart of the MVAE-DFDPnet pipeline. This flowchart elucidates the three key stages: **I. Data preprocessing** involves the integration of diverse sources of drug and protein-related information to construct unique, heterogeneous interaction networks for each view; the creation of a PCO matrix using a random surfing model, capturing the intricate network topological information of the drugs and proteins; Subsequent conversion of PCO into PPMI matrix. **II.MVAE** embeds each column of drug or protein features into a low-dimensional vector that retains the ability to reconstruct the original topological properties present within every interaction network. **III. CDF** infers a DPI as a binary value based on the concatenated learned embedding of the drug-protein pair.

#### 1) Data preprocessing

Our model utilizes heterogeneous biological networks as inputs, which are subjected to a standardized preprocessing routine independently to ensure consistency.

The random surfing model, which we employ to construct the PCO matrix, is an adaptation of the PageRank algorithm, originally developed by Google’s founders to rank webpages in search engine results [32], [33]. We have adapted this model to elucidate the interconnections among various biological entities. In a biological context, the ‘links’ might signify the interactions or associations between these entities. To construct the PCO matrix, we start by assuming a random surfer navigating the biological network. The ‘surfer’ randomly moves from one entity to another, following the edges or ‘links’. The probability that the surfer ends up on a particular node or entity is calculated. This process is repeated for all entities in the network, resulting in the PCO matrix. Subsequently, we convert the PCO matrix into a PPMI matrix, a technique widely employed in natural language processing and other domains to delineate semantic correlations among elements such as words, or in our context, biological entities like genes, proteins, or diseases, within a network. The PPMI matrix, informed by the PCO matrix, emerges as a potent instrument for elucidating complex, non-linear interdependencies and sparse associations among biological entities. Lastly, we apply matrix decomposition to the PPMI matrix, deriving novel, lower-dimensional representations of the network. This step may reveal concealed structures or patterns, thus offering fresh perspectives on the biological system under investigation. The details of data preprocessing steps are described as below:

##### (i) PCO matrix

Let *G* = (*V, E*) denote the network which contains vertices *V* and edges *E*. Suppose the vertex set *V* is sorted and has *n*_1_ biological entities from category 1 and *n*_2_ biological entities from category 2. It means the size of *V* is |*V* | = *n*_1_ + *n*_2_, and the first *n*_1_ (last *n*_2_) vertices belong categories 1(2). The edge set *E* contains undirected binary interaction information, i.e. *E* = {(*v*_*i*_, *v*_*j*_)|*v*_*i*_, *v*_*j*_ ∈ *V*, ∃ a known interaction between *v*_*i*_ and *v*_*j*_}. The info of *E* can be also represented as an adjacency matrix **A** ∈ {0, 1}^|*V* |*×*|*V*|^, where the entry **A**_*i,j*_ = 1 if (*v*_*i*_, *v*_*j*_) ∈ *E* and **A**_*i,j*_ = 0 other-wise. We define the operation *g*(*·*) that transforms a network’s adjacency matrix into a PCO matrix *g*(**A**) ∈ ℝ^|*V* |*×*|*V*|^:

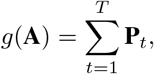

where

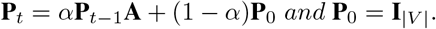

The *i*-th row of **P**_0_ is one-hot vector meaning that we start a random walk from the vertex *v*_*i*_. In each iteration process, the random surfing p rocess w ill c ontinue w ith p robability of *α*, and there is a 1-*α* probability to return to the original vertex and restart this process. So the *i*-th row of **P**_*t*_ is the probabilistic distribution of appearance at some vertex after *t* transitions starting from *v*_*i*_. Therefore, *g*(**A**) collects the cooccurrence information between vertices during the number of steps *T* in the random walk.

##### (ii) PPMI matrix

We are using the exact same formula in [19]; here we write this formula in our notations. Given a network represented by an adjacency matrix **A** for the graph **G**, where ***A***_***i***,***j***_ indicates the strength of the relationship between nodes *v*_*i*_ and *v*_*j*_, a PCO matrix *g*(**A**) is first computed, which often normalizes or thresholds the adjacency matrix to emphasize significant connections.

We define another entry-wise operation *h*(*·*) that transforms a PCO matrix *g*(**A**) into a PPMI matrix *h*(*g*(**A**)) ∈ ℝ^|*V* |*×*|*V*|^. For each pair of nodes (*v*_*i*_, *v*_*j*_) in the network, the PPMI value *h*(*g*(**A**))_*i,j*_ is calculated as the maximum of the pointwise mutual information and zero, which can be mathematically expressed as:

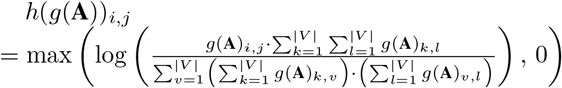

Here, the numerator represents the joint probability of observing both *v*_*i*_ and *v*_*j*_ together (normalized by their individual probabilities in the network), while the denominator corresponds to the product of their marginal probabilities. By applying the log-likelihood ratio and taking the maximum with zero, PPMI effectively removes negative associations and emphasizes stronger, more statistically significant relationships between the nodes.

##### (iii) Data integration

Now we have a PPMI matrix *h*(*g*(**A**)) retaining the network topology. The *i*-th row of matrix *h*(*g*(**A**)) contains the information of how the entity *v*_*i*_ interacts with all other entities in this network. Since we focus on only one category or part of biological entities to embed, we select the rows associated with the entities we are interested in. Let **X** denote the sub-matrix, which consists of selected rows of the PPMI matrix *h*(*g*(**A**)).

We integrate the network information for drug/protein *k* across multiple views by stacking matrices. Suppose we focus on *K*^*d*^-view data for *n*^*d*^ drugs, there are *K*^*d*^ networks 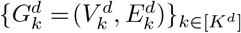 generating PPMI submatrices with selected *k k d d* rows of *n*^*d*^ drug only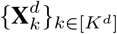. Here 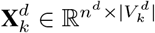, and the superscription *d* means drug. So the integrated multi-view drug-related data are represented as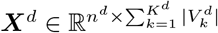 :

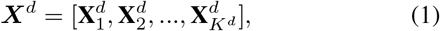

We consider the *i*^*th*^ column of ***X***^*dT*^ as the feature vector of drug *i* to be embedded by multi-view VAE, like the column framed by a red line in the Fig. 1.

Similarly, we have the superscription *p* for protein. We preprocess and then integrate *K*^*p*^-view data of *n*^*p*^ proteins into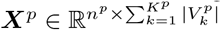 :

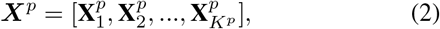

We have the *j*^*th*^ column of 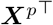 as the feature vector of protein *j* to be fed into the next module.

#### 2) MVAE for Embedding

Without loss of generality, we give as an example MVAE for drug embeddings, exactly the same model for protein embeddings. MVAE takes one preprocessed high-dimensional drug feature as input, e.g. drug *i k*=1 *k i*’s feature 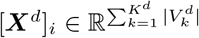 which consists of *K*^*d*^ vectors corresponding to each drug-related view. Firstly, the input feature [***X***^*d*^]_*i*_ will be embedded into a low-dimensional latent feature 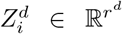 by a fully-connected neural network. Then the latent feature 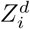 will be reconstructed back to *K*^*d*^ vectors of each view respectively by one fully-connected neural network,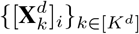. An ideal embedding should be able to recover raw input vectors of every view. So we formulate the loss function as:

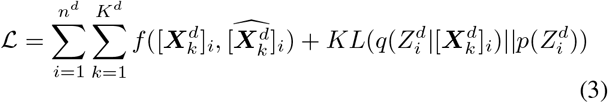

where we choose binary cross-entropy as the reconstruction loss *f*, and we use the Kullback-Leibler divergence term to regularize the difference between the learnt latent distribution and the prior distribution.

Finally, we learn a collection of drug embedding 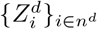 one for each drug. Similarly we have 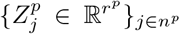 protein embedding.

#### 3) CDF for Prediction

The CDF uses ensemble and multiple-layer strategies to harness the strengths of various forest classifiers, which contribute to a powerful and robust predictive model. The diversity of the classifiers and the multi-grained scanning strategy allow the model to capture complex patterns in the data, making it suitable for a wide range of prediction tasks, including those with high-dimensional features and complex interactions.

Therefore, we use a CDF to predict DPIs. Specifically, we feed a concatenated pair embedding of drug *i* and protein *j*, i.e.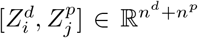, into *L* ensemble layers then output *i j* the final binary results or the score between zero and one. We include different types of binary forest classifiers to encourage diversity beyond ensembling; each ensemble layer consists of two XGBoost [34], two Random Forests [35] and two Extra Trees [36].

Each binary forest classifier outputs two non-negative values summing up as 1. Additionally, the cascade deep forest boosts the prediction performance by emphasizing the initial input, i.e. the drug-protein pair embedding. It means every layer after the first one takes all outputs of classifiers in the previous layer, along with the drug-protein pair embedding, as input. The cascade deep forest also boosts by deepening. The number of layers *N* is determined adaptively; during training, we stop adding layers when there is no noticeable decrease in the loss value. Finally, we average the outputs of the last layer to get two non-negative values summing up as 1. These two values mean the score of ‘interacting’ and ‘not interacting’ respectively.

Therefore, we take the ‘interacting or not’ result associated with the maximum score as the final binary prediction between drug *i* and protein *j*.

#### 4) Time complexity of proposed MVAE-DFDPnet

With reduced embedding dimensions, we may achieve smaller number of neurons in each layer in the time complexity of MVAE structure; while getting smaller maximum number of splits considered per feature, and smaller average depth of the tree in the CDF structure, thus improve overall model time efficiency. Detailed analysis in Supplementary materials Section A.

## III. Results

### A. Low-dimensional embedding reveals latent drug/protein families

The reconstruction loss in each view (as shown in Table II) suggests that MVAE preserves information effectively. Visualization of the low-dimensional embedding with t-distributed stochastic neighbor embedding algorithm (t-SNE) [37], [38] reveals that MVAE captures drugs/proteins information and successfully separate them into distinct clusters. This is exemplified by the 14 types of drugs by Anatomical Therapeutic Chemical (ATC)-based classification in Fig. 2a, and four classical protein families (i.e. G-protein-coupled receptors (GPCRs), kinases, nuclear receptors (NRs), ion channels (ICs), and others.) [39] in Fig. 2b. Taken together, t-SNE analysis demonstrates that the MVAE learned embedding not only greatly reduce dimensionality, but may also capture underlying biological associations within drugs and proteins.

**TABLE II:** MVAE reconstruction loss (binary Cross-entropy) across different data views.

| Drug | View | 1 | 2 | 3 | 4 | 5 | 6 | 7 | 8 | 9 |
| --- | --- | --- | --- | --- | --- | --- | --- | --- | --- | --- |
|  | Value | 0.0811 | 0.0691 | 0.0853 | 0.0317 | 0.0790 | 0.0968 | 0.0978 | 0.0903 | 0.1242 |
| Protein | View | 1 | 2 | 3 | 4 | 5 | 6 |  |  |  |
|  | Value | 0.0616 | 0.0417 | 0.0192 | 0.0561 | 0.0606 | 0.0586 |  |  |  |

**Fig. 2:**
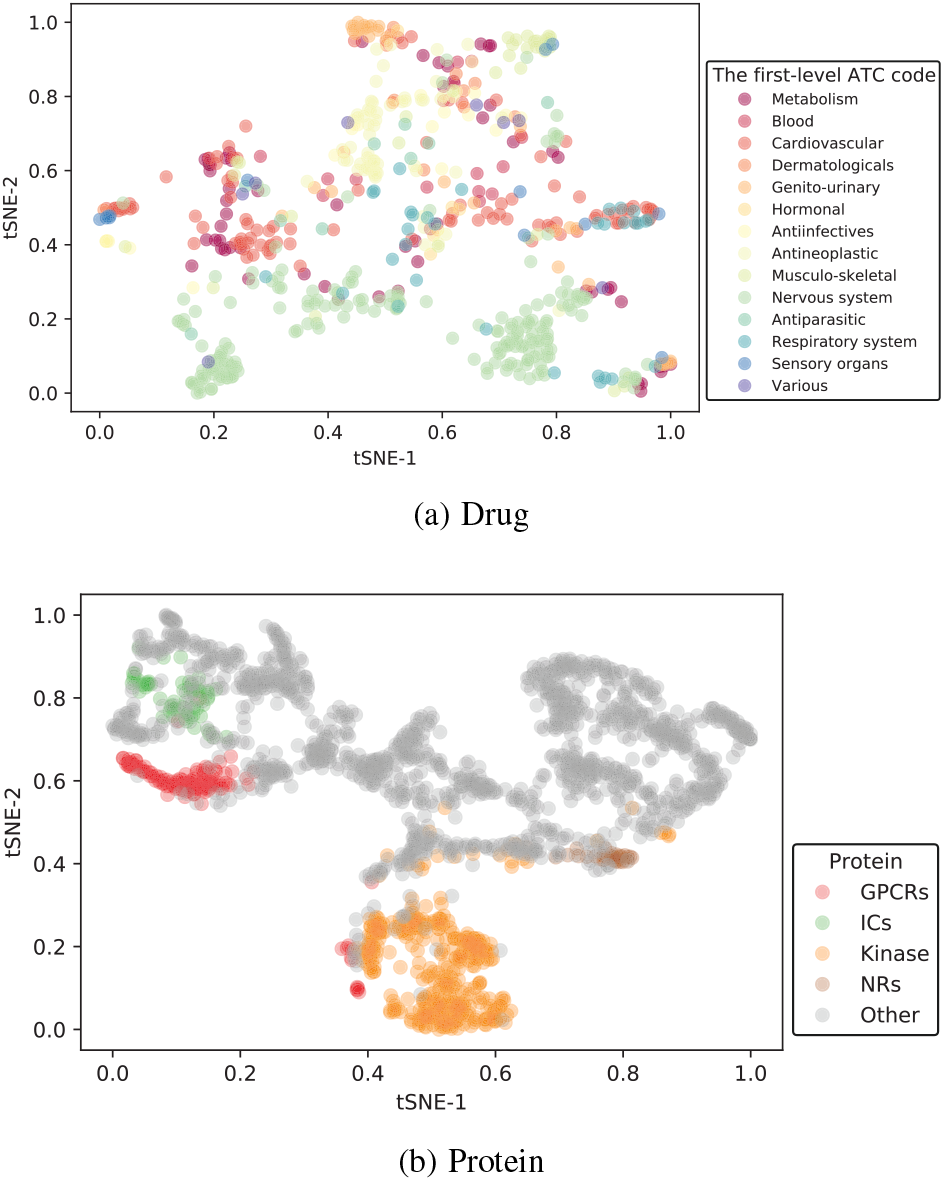
Visualization of the learned drug and protein embedding via t-SNE [37], [38]. The visualizations were created based on embeddings learned through MVAE-DFDPnet. **(a)** Drugs color-coded according to the first level of the ATC-based classification (http://www.whocc.no/atc/). **(b)** Proteins color-coded by their corresponding drug target families.

### B. MVAE-DFDPnet outperforms baseline methods

We compare MVAE-DFDPnet with the following baseline methods:

1. KBMF2K uses kernelized Bayesian matrix factorization with twin kernels for prediction [40].
2. DTINet learns low-dimensional vector representations from heterogeneous data and then applies inductive matrix completion for prediction [5].
3. NeoDTI utilizes the neighborhood information of the network for prediction [17].
4. deepDTnet obtains low-dimensional vector representations auto-encoder algorithm by and utilizes positive-unlabeled matrix completion algorithm for prediction [19]
5. AOPEDF uses an arbitrary-order proximity and a cascade deep forest classifier to infer new interactions [20].

We test different hyper parameters (Supplementary Table S1 and S2) of MVAE and evaluate the MVAE-DFDPnet model performance using the area under ROC curve (AUROC) and the area under the precision-recall curve (AUPR). Since negative samples are many more compared to positive samples in our data, we randomly sampled the negative samples to reach ratio of 1:1 in each test. All methods are performed 10 times of random 5-fold cross-validation and computed the average performance. Fig. 3 shows the average AUROC and AUPR for each method. The data indicate that MVAE-DFDPnet surpasses leading-edge methods such as KBMF2K, DTINet, NeoDTI, deepDTnet, and AOPEDF. Notably, MVAE-DFDPnet achieves impressive results with drug-protein embeddings of merely four dimensions (AUROC = 0.973 and AUPR = 0.974, as shown in Table III using a pairwise train-test split), outstripping most prior methods. As the dimensionality of the embeddings increases to 200 and 2,000, MVAE-DFDPnets performance further enhances, yielding an accuracy on par with the top-performing method, AOPEDF (AUROC = 0.975 and AUPR = 0.974), while utilizing a significantly reduced dimensional space for embeddings (200 compared to AOPEDFs 1,650). The result of compared AOPEDF with different dimensions is in Supplementary Table S3.

**TABLE III:** Evaluation performance of MVAE-DFDPnet compared with state-of-art methods. The associated variance is in parentheses.

| Method | Embedding Dimension | AUROC | AUPR |
| --- | --- | --- | --- |
| KBMF2K | 200 | 0.937 | 0.946 |
| DTINet | 500 | 0.932 | 0.943 |
| NeoDTI | 2048 | 0.971 | 0.97 |
| deepDTnet | 2000 | 0.963 | 0.974 |
| AOPEDF | 1650 | 0.975 | 0.974 |
| MVAE-DFDPnet | 4 | 0.975(0.008) | 0.976(0.008) |
|  | 20 | 0.982(0.006) | 0.984(0.008) |
|  | 200 | 0.985(0.006) | 0.986(0.008) |
|  | 2000 | **0.986(0.006)** | **0.986(0.006)** |

**Fig. 3:**
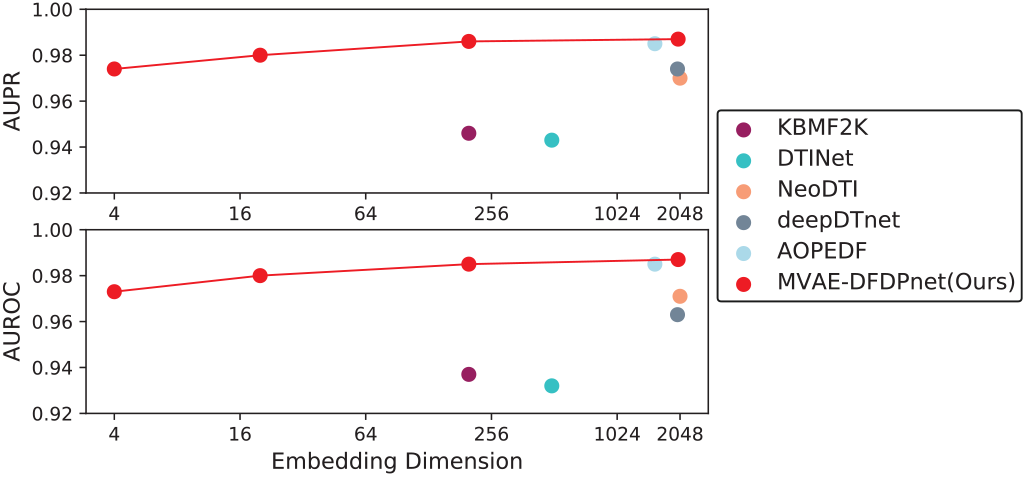
Performance comparison of MVAE-DFDPnet against State-of-the-art methods at various embedding dimensions.

### C. Robustness and generalizability of MVAE-DFDPnet

Random splitting of drug-protein pairs for testing may result in the inclusion of drugs or proteins in the test set that have been seen during training, potentially leading to an overestimation of the model’s performance, and overly optimistic conclusions. To address this issue, we adjust sample splitting to test the MVAE-DFDPnet method on entirely novel drugs, novel proteins, or both novel scenarios (Fig. 4a). The AUROC and AUPR remain at 0.9 even when both the protein and the drug in the test set are entirely new (Fig. 4b). This demonstrates the robustness of MVAE-DFDPnet in predicting DPIs for new drugs or proteins, even without prior knowledge of the drug or protein.

**Fig. 4:**
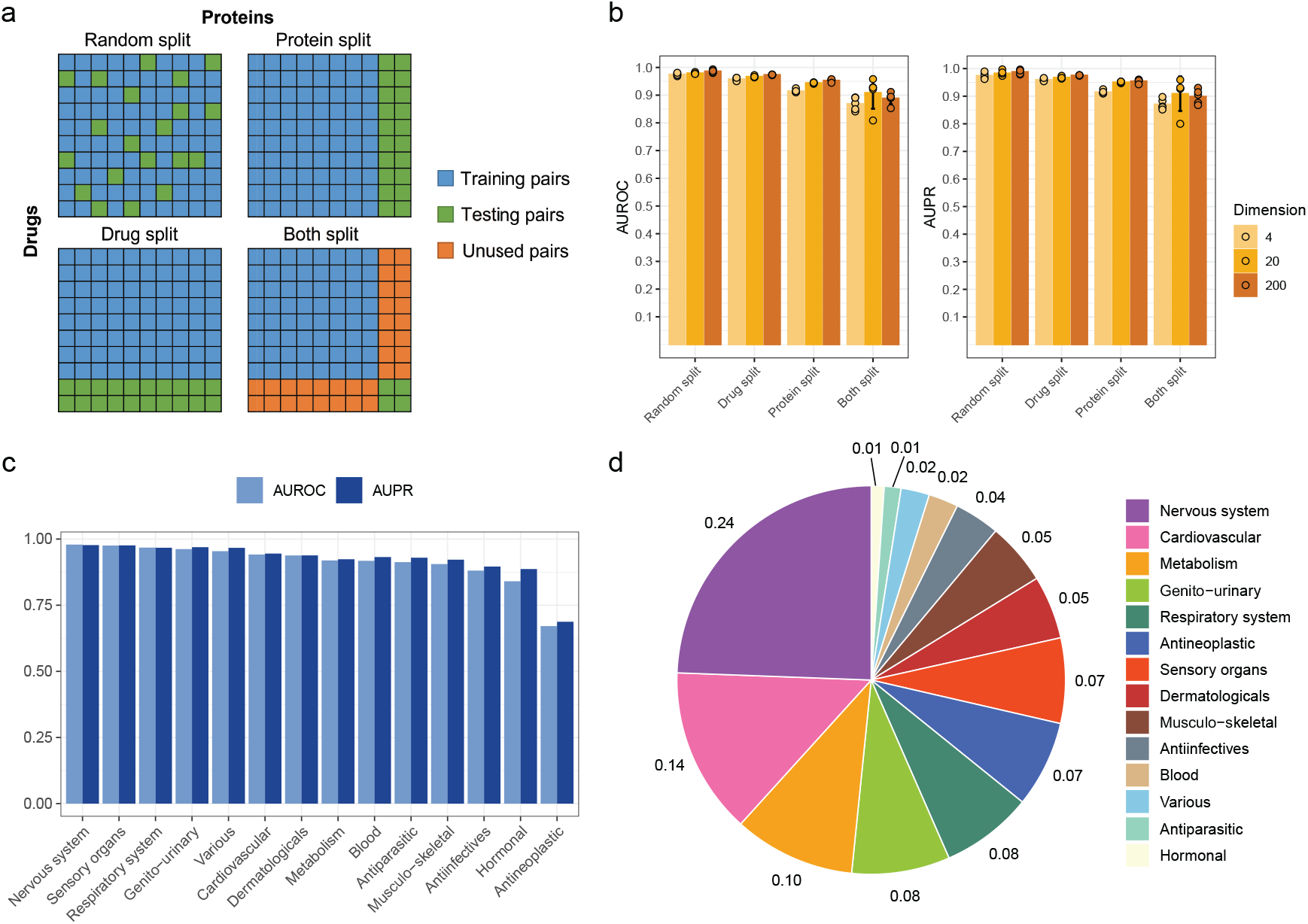
The performance evaluation results of MVAE-DFDPnet on several challenging scenarios. (a) The schematic diagram of splitting drug-protein pairs into training and testing sets. (b) The performance of MVAE-DFDPnet to accurately predict the drug-protein interaction of completely novel drugs, proteins, or both novel. (c) The performance of the MVAE-DFDPnet in predicting novel drugs of ATC-based classifications. (d) The proportion of drugs in each ATC classification in this study.

Further, drugs belonging to the same class might be structurally and functionally similar, therefore excluding an entire drug class in training data would pose an even more challenging task for DPI prediction. We test MVAE-DFDPnet in predicting DPIs of each ATC drug class while excluding the given class of drugs from training data (Fig. 4c), resulting in most ATC class AUROC and AUPR scores of *>*0.9 indicating good generalizability across most drug class. Note that the absence of antineoplastic drugs in training data has a significant impact on DPI prediction. The number of drugs in each ATC drug class (Fig. 4d) is not associated with their impact on the DPI prediction.

### D. MVAE-DFDPnet reveals novel DPIs

We validate novel DPIs with DGIdb database [41]; DGIdb supports 1,705 known + 1,379 novel DPIs. Among top 60 DGIdb-validated predictions in Supplementary Table S4, we find that many of them can also be supported by the previously known experimental or clinical evidence in the literature [42]– [51].

We visualize the top 100 novel DPIs between 30 drugs and 40 proteins. As shown in Fig. 5, kinases and their drug interactions tend to form a cluster that is separated from those of GPCR and others. In the kinase network, the interactions are between multiple tyrosine kinase inhibitors and kinases, while the majority of the interactions are supported by previously reported experimental evidence in the literature. For example, Sunitinib is a multi-target tyrosine kinase inhibitor, indicated for the treatment of renal cell carcinoma and imatinib-resistant gastrointestinal stromal tumor. LIMK1 is an important member of LIMK/cofilin signaling and participates in actin reorganization, cell migration, and tube formation. One research found that the indolin-2-one derivatives potently inhibit the LIMK1/cofilin signaling pathways, while sunitinib is a representative drug that emerged from indolin-2-one [52]. The prediction between Sunitinib and LIMK1 merit further studies, which may point to therapy that target cancer invasive behavior. Similarly, EGFR-mutant non-small cell lung cancer is known to respond to EGFR inhibitors such as Gefitinib. A study identified several genetic determinants of EGFR TKI sensitivity through genome-wide CRISPR-Cas9 screening. Researchers found that sgRNA targeting PKN2 strongly sensitized HCC827 cancer cells to gefitinib treatment. Another synergic gene RIC8A was found attenuating YAP signaling, which might be modulated via both LATS1/2 [53]. This demonstrates the potential insights that can be gained from our novel DPI predictions.

**Fig. 5:**
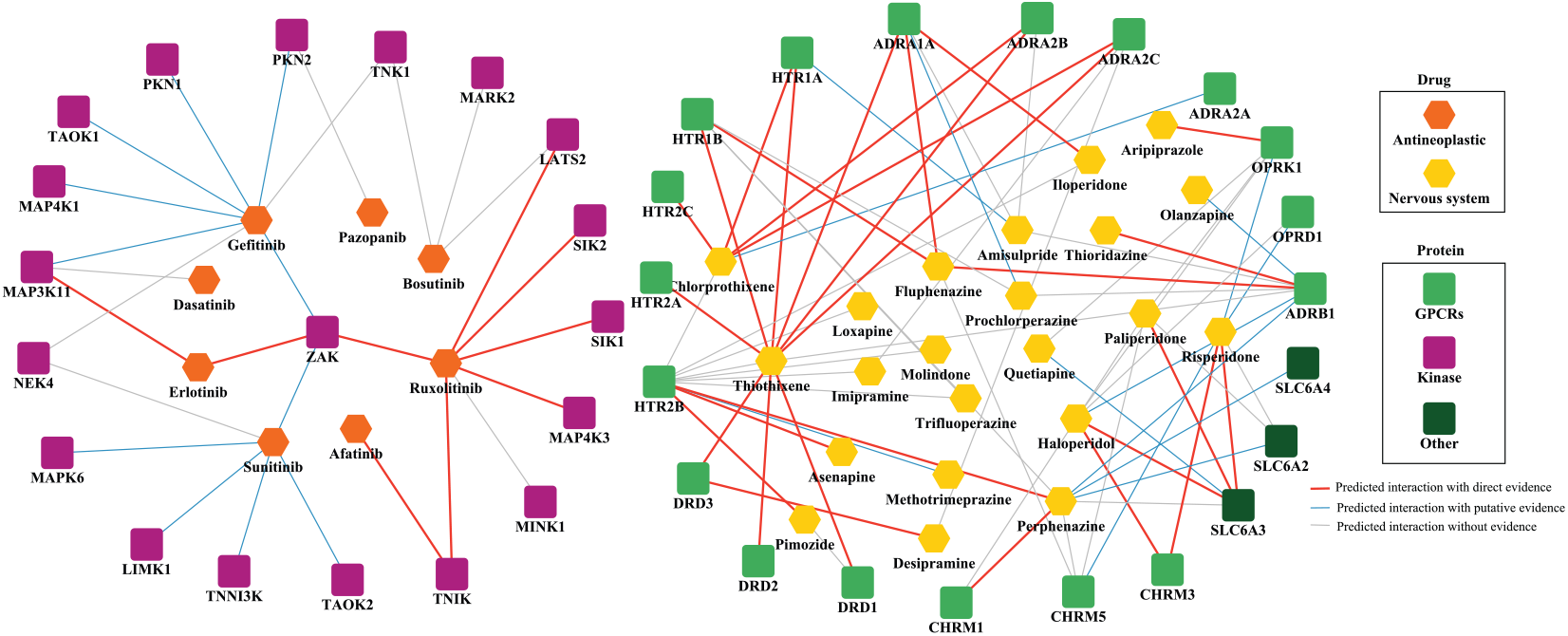
Top 100 novel DPIs predicted by MVAE-DFDPnet. This figure showcases the top 100 novel DPIs as predicted by our MVAE-DFDPnet model. The drugs are grouped by the first ATC-level code, in this case, Antineoplastic and Nervous system. Many novel DPI predictions are corroborated by experimental data, database entries, or clinical findings reported in the literature. DPIs with direct evidence available are marked by red lines, those with putative evidence available are marked by blue lines; while DPIs lacking such evidence are represented by grey lines.

### E. Case study: antiepileptic drugs

We attempt to focus on the drugs in our dataset that fall under the ATC classification system N03A (Nervous system/antiepileptics). Antiepileptic drugs (AEDs) are typically prescribed for chronic, long-term use in patients with epilepsy, often extending over several years [54]. The prediction scores for these drugs are generally lower than those of the top 100 interactions previously discussed. Our MVAE-DFDPnet model was able to computationally identify 440 potential interactions linking 117 proteins with 12 AEDs. We further illustrated the top 100 interactions encompassing 10 AEDs and 36 proteins in Fig. 6. The potential for these predicted interactions to be supported by existing research seems to correlate positively with their prediction scores. It is noteworthy that ion channel modulators such as phenytoin, clonazepam, and vigabatrin exhibited unique interaction profiles, indicating diverse mechanisms of action. Furthermore, our model predicted a wide range of interactions between most of the AEDs and an array of GPCRs. This prediction could shed light on the ongoing discussion around the therapeutic potential of GPCRs for treating acquired epilepsy [55]. Thus, these findings could guide the development of new AEDs or therapeutic strategies and be utilized to repurpose existing drugs for acquired epilepsy treatment, potentially accelerating the drug development process.

**Fig. 6:**
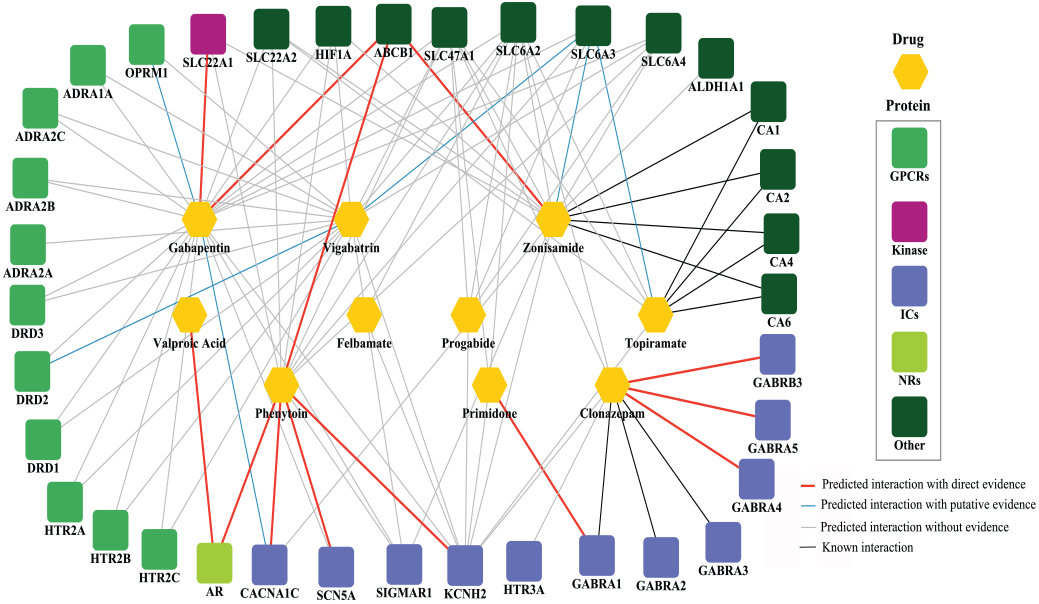
Antiepileptic drug (AED)-related DPI network featuring the top 100 novel DPIs predicted by MVAE-DFDPnet. Drugs classified under the N03A antiepileptics category are represented by yellow hexagons. Proteins are depicted as squares, each color-coded according to their respective protein families. DPIs are differentiated by colored lines, each of which represents the type or availability of supporting evidence.

## IV. Conclusion and Future Work

In this study, we present a novel deep learning architecture, MVAE-DFDPnet, tailored for DPIs prediction. This framework integrates a multi-view variational autoencoder with a cascading deep forest classifier, offering a refined and potent method for inferring novel DPIs.Our experimental results demonstrate the superiority of MVAE-DFDPnet in DPI prediction, outperforming existing state-of-the-art techniques while utilizing a significantly reduced dimensionality of drug-protein embeddings, with good generability and robustness. Future endeavors may include the integration of additional biological data sources to augment the model’s input dataset.Further optimization and enhancement of the MVAE-DFDPnet components could yield even more accurate and efficient DPI predictions. Lastly, empirical validation of the DPI predictions in laboratory settings would reinforce the practical and scientific merits of our approach.

