## supplement for "Redefining the Game: MVAE-DFDPnet’s Low-Dimensional Embeddings for Superior Drug-Protein Interaction Predictions"

Table S1. The chosen architecture of each network in MAVE-DFDTnet.

| **Class** | **Encoder** | **Decoder(View ID)** |
| --- | --- | --- |
| Drug | 732*9 500 200 100 **d** | 100 200 500 732 (1) |
|  |  | 100 200 500 732 (2) |
|  |  | 100 200 500 732 (3) |
|  |  | 100 200 500 732 (4) |
|  |  | 100 200 500 732 (5) |
|  |  | 100 200 500 732 (6) |
|  |  | 100 200 500 732 (7) |
|  |  | 100 200 500 732 (8) |
|  |  | 100 200 500 732 (9) |
| Protein | 1915*6 500 200 100 **d** | 100 200 500 1915 (1) |
|  |  | 100 200 500 1915 (2) |
|  |  | 100 200 500 1915 (3) |
|  |  | 100 200 500 1915 (4) |
|  |  | 100 200 500 1915 (5) |
|  |  | 100 200 500 1915 (6) |

Note: In this study, we set ω=0.98, and T= 3 (the number of RW steps) in the random surfing step of MVAE-DFDTnet. We then chose the parameter of MVAE and deep forest. Specifically, we first fixed the embedding dimension **d** = 2/10/100/1000 according to previous experience. Then we designed different architectures of MVAE-DFDTnet for 15 networks separately. We varied the learning rate, dropout rate, NN layers and NN hidden units of each network and decide the architecture of each network one by one according to the prediction results indicated by 5-fold cross validation

Table S2. Experimental parameters.

| **Parameter** | **Value** |
| --- | --- |
| Epochs | 500 |
| Batch size | 100 |
| Optimizer | adam |

Table-S3: Evaluation performance between MVAE-DFDTnet and AOPEDF.

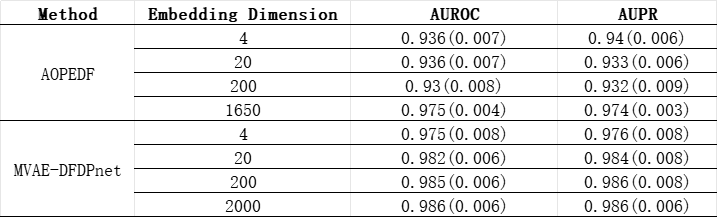

Table-S4:MVAE-DFDPnet predicts 60 novel DPIs validated by evidence in DGIdb database [41]. The score cutoff for the top 60 novel interactions with DGIdb evidence is 0.999. For all interactions above this cutoff, the false positive rate is 35.75%; 60 out of the top 1073 false positive interactions can be validated with evidence.

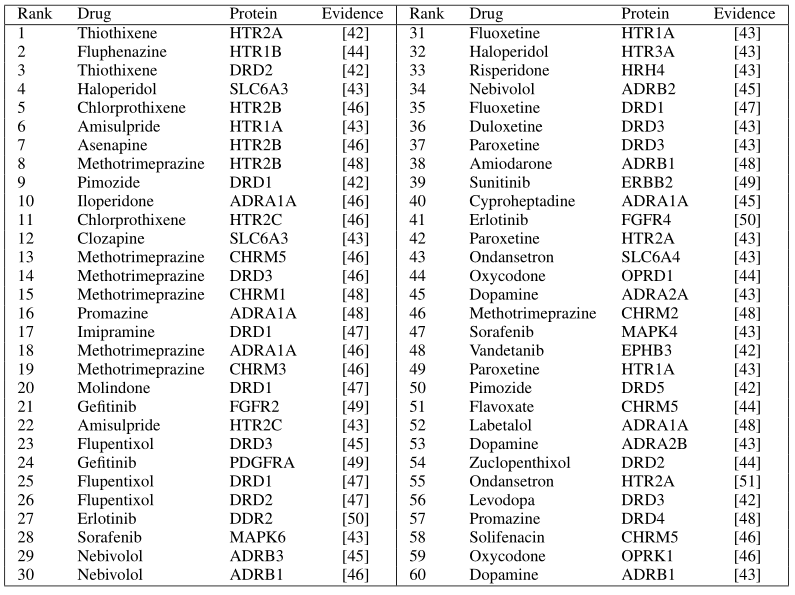

**Section A(Time complexity)**：

The time complexity of various components in a MVAE and CDF can be analyzed separately. MVAE consists of two main parts: the encoder and decoder networks, each applied to multiple views or sources of data.

- Encoder: For each view, the neural network typically involves layers with operations like matrix multiplications, activation functions, and pooling. Assuming there are
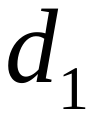
input features per view and
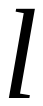
 hidden layers with
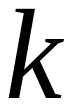
 neurons in each layer for one view’s encoder, the time complexity is approximately
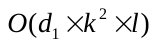
. Since we have multiple views, the overall complexity would be
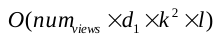
.
- Decoder: The decoder architecture is similar to the encoder but in reverse, so its time complexity would also be
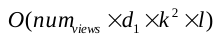
.
- Variational Inference: This process involves sampling from the learned distribution, calculating the KL divergence, and optimizing the model using backpropagation. With batch size
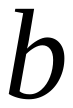
, this step has a time complexity of
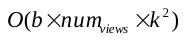

Overall, MVAE's time complexity would be dominated by the neural network layers, which is roughly
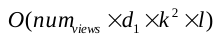
.

CDF is an ensemble learning method that combines multiple decision trees(RandomForest, XGBoost, and ExtraTrees). Here’s the time complexity breakdown:

- Training individual Decision Tree: At each node, it performs best split search over all possible feature-value combinations, which has a time complexity of
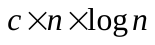
, where
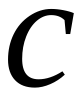
 is the maximum number of splits considered per feature, and
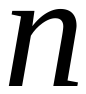
 is the number of samples.
- Building the tree: Recursively splitting nodes until a stopping criterion is met (like minimum sample size or maximum depth), which incurs an additional
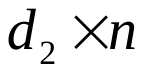
, where
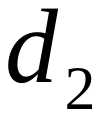
 is the average depth of the tree.For a cascade of
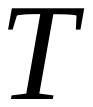
trees, the time complexity for training would be
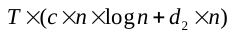
.
- Prediction: Each tree makes a prediction independently, requiring a traversal through the tree for each instance. Time complexity is
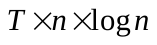
.
- Therefore, the overall time complexity for a CDF with XGBoost, Random Forest, and Extra Trees is
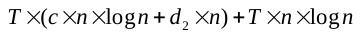
.

Taken together, with reduced embedding dimensions, we may achieve smaller
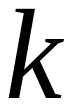
 in the equation of time complexity of MVAE structure; while getting smaller
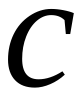
and
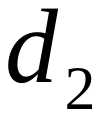
 in the time complexity of CDF structure, thus improved overall model time efficiency.
